# Label-free assessment of pre-implantation embryo quality by the Fluorescence Lifetime Imaging Microscopy (FLIM)-phasor approach

**DOI:** 10.1101/286682

**Authors:** Ning Ma, Nabora Reyes de Mochel, Paula Duyen Anh Pham, Tae Yeon Yoo, Ken WY. Cho, Michelle A. Digman

**Affiliations:** Department of Biomedical Engineering, University of California, Irvine, CA; Laboratory of Fluorescence Dynamics (LFD); Department of Developmental and Cell Biology, University of California, Irvine, CA

## Abstract

Development of quantitative, safe and rapid techniques for assessing embryo quality provides significant advances in Assisted Reproductive Technologies (ART). We apply the phasor-FLIM method to capture endogenous fluorescent biomarkers of pre-implantation embryos as a non-morphological caliber for embryo quality. Here, we identify the developmental, or “D-trajectory”, that consists of fluorescence lifetime from different stages of mouse pre-implantation embryos. The D-trajectory correlates with intrinsic fluorescent species from a distinctive energy metabolism and oxidized lipids, as seen with Third Harmonic Generation (THG) that changes over time. In addition, we have defined an Embryo Viability Index (EVI) to distinguish pre-implantation embryo quality using the Distance Analysis, a machine learning algorithm to process the fluorescence lifetime distribution patterns. We show that the phasor-FLIM approach provides a much-needed non-invasive quantitative technology for identifying healthy embryos at the early compaction stage with 86% accuracy. This may increase embryo implantation success for *in vitro* fertilization clinics.

**Highlights:**

- A label-free method of tracking metabolic trajectories during pre-implantation mouse embryo development.
- A non-invasive approach for assessing embryo quality and viability by a phasor-FLIM analysis.

## Introduction

Determining embryo quality during *in vitro* fertilization (IVF) is one of the most important steps toward successful pregnancy^1^. The standard non-invasive method to assess embryo quality and viability relies on the visual inspection of embryo morphology according to predefined criteria such as cell division patterns, the number of pronucleoli in cleavage stages^2,3^, and the physical characteristics of the blastocyst^4^. Assisted reproduction through morphological evaluation is labor intensive and highly dependent on the performance of individual physicians trained in these techniques. Development of more quantitative and objective means for assessing embryo quality that are simpler, safer, and faster could provide significant advantages in assisted reproduction by enabling single embryo transfers rather than the implantation of multiple embryos in order to increase the likelihood of a successful pregnancy.

Given the limitations of morphological evaluation, several technologies have been explored for the assessment of embryo viability. These include the measurement of metabolites in embryonic culture media, as well as genomic and proteomic profiling of the embryos themselves^5^. For example, spectroscopic approaches have been utilized to measure the amount of metabolites such as pyruvate, lactate, and glucose in the media during embryo culture^6–8^. However, these approaches are time-consuming and require highly-trained personnel to analyze the complex data^9^. Both genomic and proteomic profiling are equally time consuming and can cause damage to the embryo during the procedure.

Here, we apply the phasor-fluorescence lifetime imaging microscopy (FLIM) method and examine the dynamic endogenous biomarker (metabolites as described below) changes during preimplantation embryo development. Based on the quantifiable physiological property changes, we correlate the biomarker changes to the embryo viability (Fig. 1). This non-invasive phasor-FLIM analysis is sensitive, quick and intuitive.

**Figure 1:**
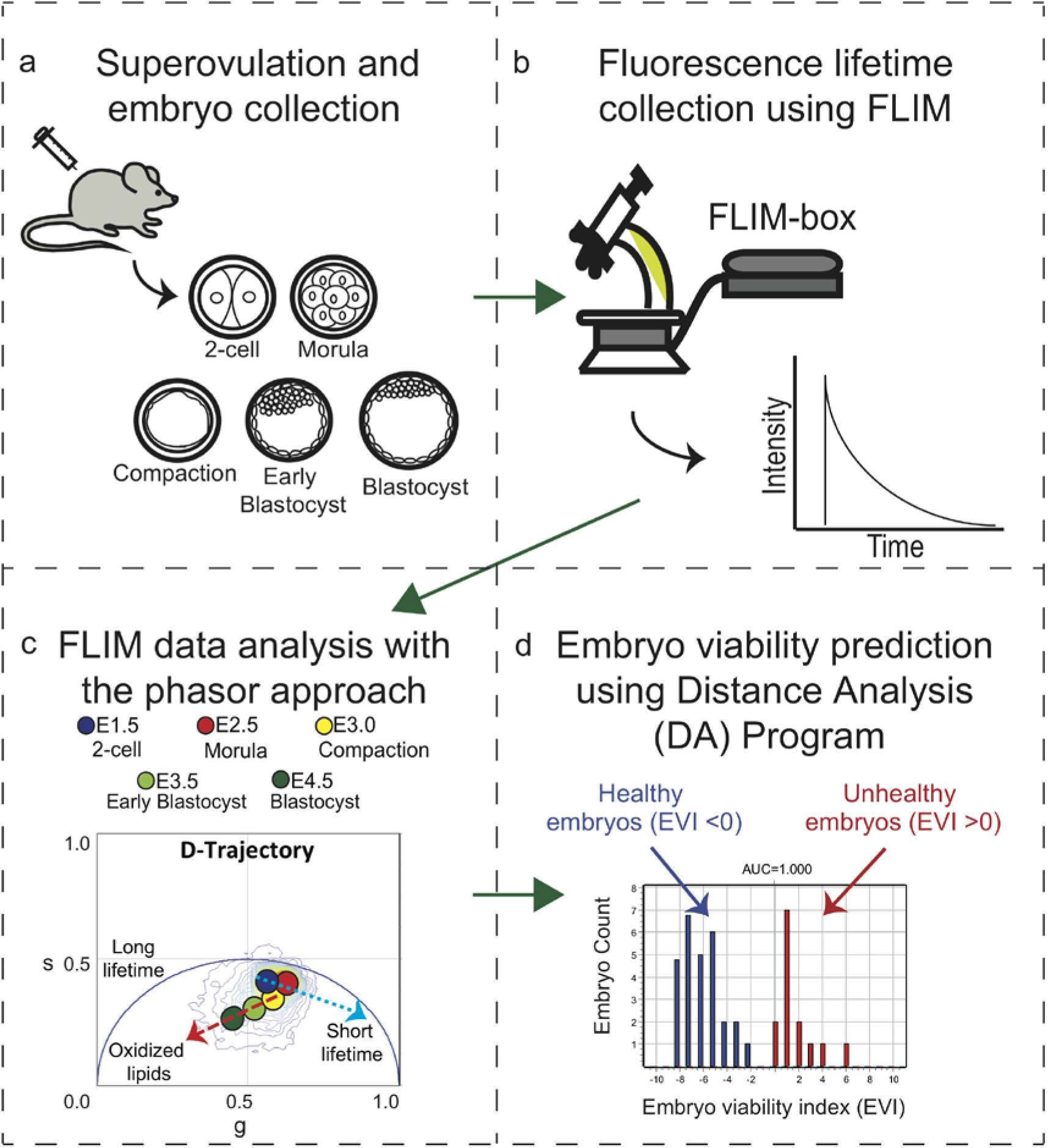
Schematic of the workflow of the experimental design. a) We collected FLIM images of embryos from superovulated female mice at the following developmental stages: 2-cell, morula, compaction, early blastocyst, and blastocyst. b) Intrinsic fluorescence lifetimes for each embryo are collected using a Zeiss 710 microscope coupled with a FLIM-box. c) The FLIM data analysis of the pre-implantation mouse embryo development was performed using the phasor approach. d) Distance Analysis (DA) program was applied to predict embryo viability.

FLIM produces an image, based on the exponential decay rates at each pixel from a fluorescent sample. The fluorescence lifetime of the fluorophore signal is measured to create the image via FLIM^10^ (Supplementary Fig. 1a). When FLIM is coupled with two-photon excitation microscopy, molecules are excited at longer wavelengths (with lower energy photons). This prevents photo damage and allows deeper imaging, resulting in superior image quality^11^. Since endogenous molecules such as collagen, retinoids, flavins, folate and NADH (nicotinamide adenine dinucleotide) are fluorescent in live cells^12–14^, we can collect fluorescence lifetime data to identify these intrinsic fluorescent species. The contributions from these different biochemical species are indicators of an embryo’s biochemical property^15,16^. In our approach, we measure the fluorescent lifetime signal from integrated images acquired and transform the raw data using the Fourier transformation to the average arrival time of emitted photons in each pixel, represented by polar coordinates “g” and “s” in the transformation function^13^ (Fig. 1c, Supplementary Fig. 1a). This allows us to present the data in a two-dimensional graphical representation of the lifetime distributions, known as the phasor plot, for each pixel in the FLIM image (Supplementary Fig. 1).

Here we have applied the phasor-FLIM approach to pre-implantation mouse embryos and have captured detailed data on their metabolic states at various developmental stages. At each stage, the mouse embryo displays a characteristic phasor-FLIM signature. For the first time, we defined a unique graphical metabolic trajectory that correlates with energy metabolism and embryo development, which we call the developmental trajectory or “D-trajectory”. Initially embryos uptake pyruvate during glycolysis as their main energy source^17^. As the embryos develops to later stages, the need for ATP increases in order to activate transcription for proliferation. Then, the embryos switch from glycolysis to oxidative phosphorylation, primarily using glucose as their energy source, which also changes the relative redox potential (NAD+: NADH ratio)^18^. The spectroscopic signatures from each of these changes are detected and can be used as criteria to identify healthy embryos at each stage in development. We find that the D-trajectory of pre-implantation embryos cultured in nutrient-deficient media deviates significantly from that of the normal media, indicating that lifetime trajectories can be used to detect metabolic alterations in embryos. We have identified a combination of mathematical parameters that are statistically different between healthy and unhealthy pre-implantation embryos based on machine learning information. Therefore, the phasor-FLIM approach provides an objective, non-invasive, and quantitative method to assess the quality of mammalian embryos.

## Results

### The lifetime D-trajectory of pre-implantation embryos

Two different mouse strains (a non-inbred CD1 and an inbred C57BL/6NCrl) were used to acquire a comprehensive representation of the phasor-FLIM distribution patterns of embryos during pre-implantation development (Fig. 2 and Supplementary Fig. 2a). Fluorescent lifetimes of endogenous fluorescent species, excited at 740nm, were collected at the 2-cell (E1.5), morula (E2.5), compaction (E3.0), early blastocyst (E3.5) and blastocyst stage (E4.5), and pseudo-colored according to the phasor coordinates (Fig. 2a, b). The phasor coordinates, which is the averaged fluorescent lifetime, of the 2-cell and morula stage embryos have a unique lifetime distribution pattern distinct from all other cell and tissue types measured (blue arrow, Fig. 2b)^19–22^. This unique phasor lifetime position may reflect special characteristics of totipotent cells, which mirror low oxygen consumption and preferential utilization of pyruvate oxidation^23^. On the other hand, compaction to blastocyst stages display average phasor coordinates typically observed in pluripotent cells (red arrow, Fig. 2b)^20,22^. We refer to this characteristic developmental time course lifetime distribution pattern as the developmental trajectory or “D-trajectory”. Phasor-FLIM lifetime distributions of individual embryos from both outbred and inbred mouse strains (Fig. 2c, d) follow the similar developmental trend D-trajectory. In order to examine whether genetic background of mice influences the D-trajectory, we compared the trajectories of both CD1 and C57BL/6NCrl strains (Fig. 2c, d). While the average lifetimes (g and s values) at specific embryonic stages are somewhat variable, the overall D-trajectory distribution (blue and red arrows) of C57BL/6NCrl is similar to that of CD1 mice. We conclude that the D-trajectory is a characteristic distribution behavior observed among pre-implantation mouse embryos. Lastly, we have applied time-lapse FLIM imaging to individual embryos (n=16), and continuously followed at 3-hour time intervals from 2-cell (E1.5) to blastocyst stage (E4.5) for approximately 60 hours. The *in vitro* developmental trajectory (Supplementary Fig. 2c) of each embryo mirrors the D-trajectory (Fig. 2b, with blue and red arrows). In sum, the combined two lifetime trajectories (blue and red arrows) encompass the overall D-trajectory for normal pre-implantation embryo development.

**Figure 2:**
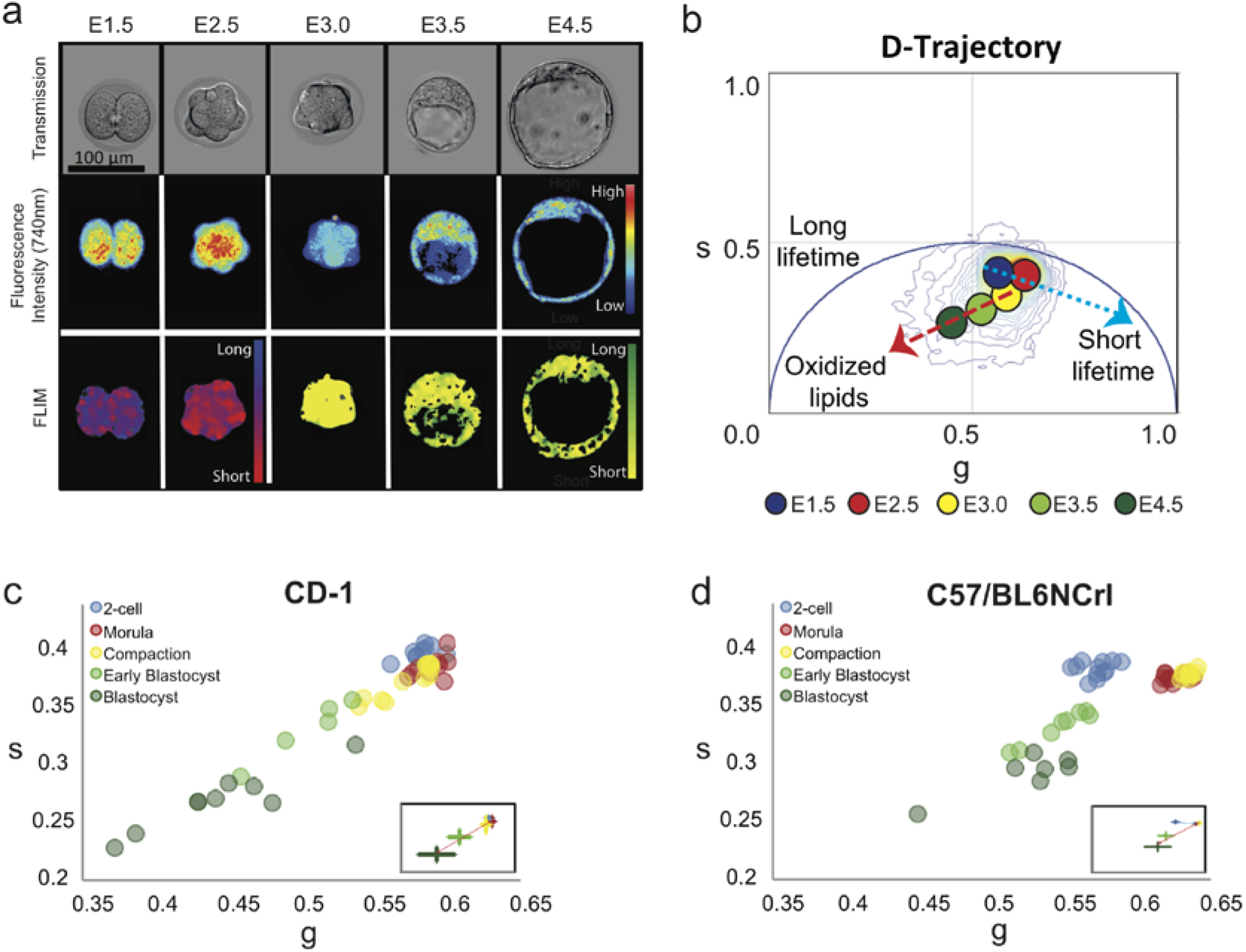
The lifetime trajectory of pre-implantation embryos correlates with embryonic development. a) Transmission (top row), fluorescence intensity (middle row, 740nm excitation) and FLIM (bottom row) images of representative pre-implantation CD1 mouse embryos at 2-cell, morula, compaction, early blastocyst, and blastocyst stage. In the FLIM images, the pseudo color displays the fluorescence lifetime. b) Phasor-plot of average fluorescence lifetime of CD1 embryos at the indicated developmental stages demonstrating the D-trajectory (D for development). A blue arrow indicates the fluorescence lifetime change from E1.5 to E2.5 and a red arrow shows the change from E3.0 to E4.5. c-d) Scatter plots show the D-trajectory for CD1 and C57BL/6NCrl embryos. Small window shows the average and standard deviation of each stage. CD1: 2-cell (n=29), morula (n=11), compaction (n=33), early blastocyst (n=50) and blastocyst stage (n=35); C57BL/6NCrl: 2-cell (n=25), morula (n=22), compaction (n=21), early blastocyst (n=38) and blastocyst stage (n=42). c) D-trajectory of CD1 embryos (2-cell, n=8; morula, n=8; compaction, n=12; early blastocyst, n=5; blastocyst, n=8. and d) D-trajectory of C57BL/6NCrl embryos (2-cell, n=7; morula, n=3; compaction, n=17, early blastocyst, n=8; blastocyst, n=21). N= number of embryos analyzed.

Reactive oxygen species (ROS) plays a key role in cellular metabolism and homeostasis^24,25^ and ROS production has been linked to an increase in oxidized lipids^26,27^. The red arrow (the right to left-downward shift, Fig. 2b) in the D-trajectory is presumably due to an increasing fractional contribution of ROS as well as the oxidized lipids which have a fluorescence lifetime distribution of 7.89ns and fall on the same published location (coordinates) of the semi-circle in the phasor plot (Supplementary Fig. 1b)^13^. This behavior is consistent with the model that an increase in aerobic respiration and metabolism as well as β-oxidation during pre-implantation mouse development^13^ requires more efficient energy production from oxidative phosphorylation^28,29^. We have confirmed the presence of active ROS production with fluorogenic marker 2’, 7’-dichlorofluorescin diacetate (DCF-DA, also known as H_2_DCFDA) staining (Supplementary Fig. 2b).

In order to better characterize the lipid droplets distribution during embryonic development, we have employed third-harmonic generation (THG) microscopy imaging (Fig.3) with Deep Imaging Via Emission Recovery (DIVER) microscope (Fig.3). The interfaces heterogeneity can be detected with the third order nonlinearity χ^3^. Given that the process is ultra-fast for structures with THG signals, the lifetime is approximately zero. Figure 3a shows the representative THG intensity images acquired in the same field of view as that of the FLIM images of Figure 3b. The phasor plot of the THG images appear at the coordinate of s = 0 and g = 1. The 3D structure (Supplementary Movie 1a-e) of the lipid droplets of embryos from different stages^14^. Furthermore, we quantify the co-localization correlation of the long lifetime specie in the FLIM images (red) with the lipid droplets (green) in THG images (Fig. 3c, d). During embryonic development, the oxidized lipid signature, color-coded in red for the long lifetime species, (same direction as red arrow in Figure 2b) accumulated. The Mander’s split co-localization correlation coefficients increase from 0.0099 to 0.3907 (where a coefficient of 1 is perfect correlation and 0 is complete lack of correlation) with embryonic development, suggesting that the phasor-FLIM distribution changes during these stages are due to increased lipid accumulation. We also characterized the lipid droplets distribution during embryonic development using the 3D THG image (Fig. 3a, e). Cleavage stage embryos have large amount of small, densely packed lipid droplets, whereas post-cleavage stage embryos have large lipid droplets of the low density. And the dramatic changes for both the lipid oxidation and lipid volume decreasing start happening after compaction stages. These findings demonstrate that the dynamic difference in lipid oxidation can be detected by phasor-FLIM.

**Figure 3:**
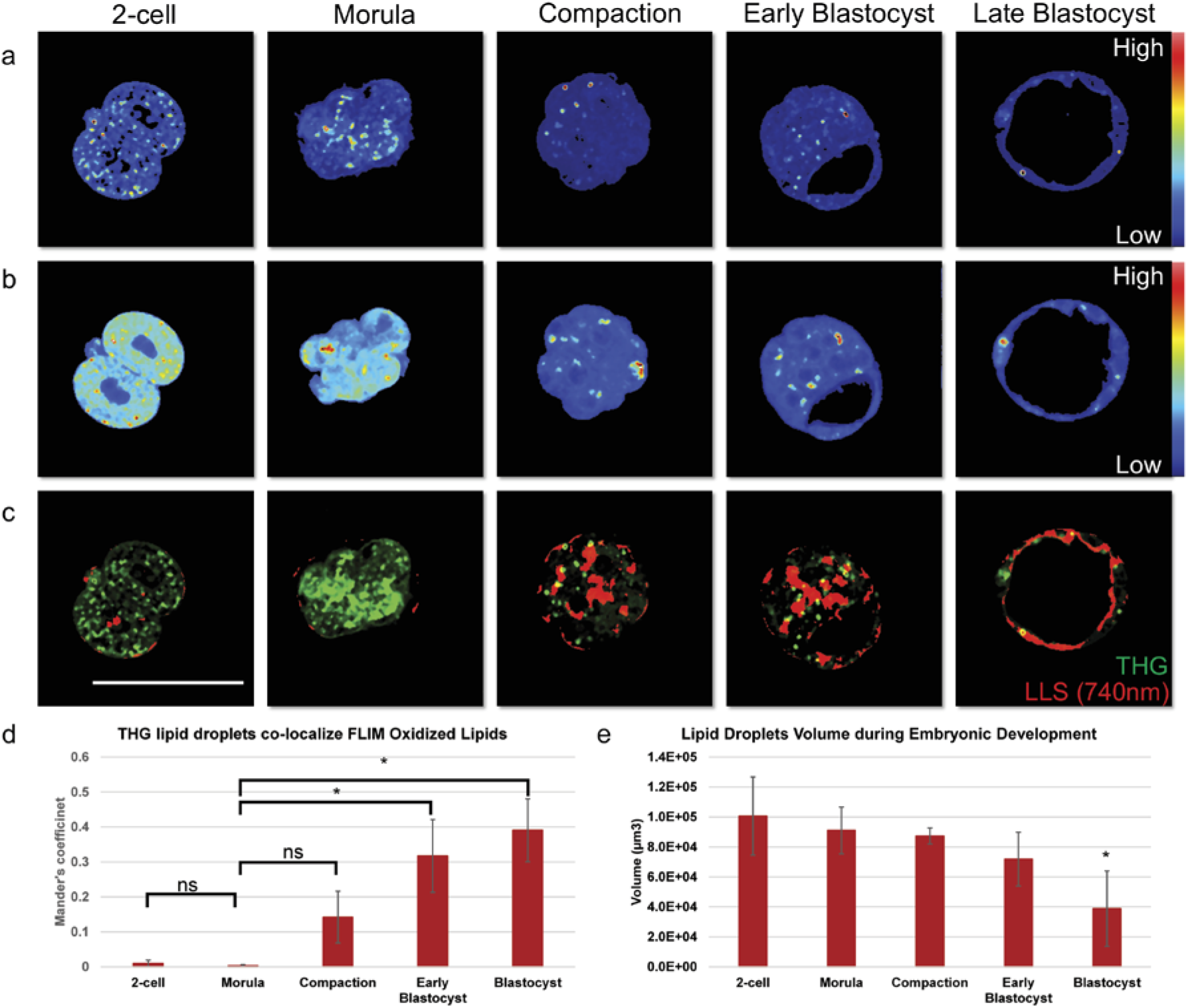
THG and intrinsic fluoresce signal show increasing oxidized lipids during embryonic development. Representative third harmonic generation images (a) and FLIM images (b) during pre-implantation embryonic development, 2-cell, morula, compaction, early blastocyst and blastocyst stage for the same field of view. From blue to red shows the intensity increase. c) Representative optical sections show co-localization (yellow) of the lipid droplet signal (green) in THG images with long lifetime species (red) in FLIM images. Scale bar sets at 100µm. d) Mander’s coefficient of the co-localization results during embryo development which shows the proportion co-localization region of the THG channel and FLIM channel correspondence with long lifetime specie-oxidized lipids. 2-cell (n=5), morula (n=3), compaction (n=3), early blastocyst (n=4) and blastocyst stage (n=3). Student t-test results (p-value) for morula to 2-cell, compaction, early blastocyst and blastocyst are 0.1923, 0.0823, 0.0091, and 0.0174 respectively. e) Lipid droplets volume characterization during pre-implantation embryo development. 2-cell (n=5), morula (n=5), compaction (n=4), early blastocyst (n=4) and blastocyst stage (n=6). Student t-test results (p-value) for morula to 2-cell, compaction, early blastocyst and blastocyst are 0.5066, 0.6367, 0.1416, and 0.0072 respectively.

### Fluorescence lifetime trajectories reveal metabolic states of preimplantation mouse embryos

The D-trajectory is complex because it is composed of lifetimes from various endogenous fluorescent biochemical species. We first hypothesized that the major components responsible for the shifts in the D-trajectory is intracellular NADH changes based on its fundamental role in energy production during embryogenesis. To test this, we first measured the metabolic activity of intracellular NADH^30,31^. The bound form of NADH is linked to energy production through oxidative phosphorylation, whereas the free form of NADH is associated with glycolysis^21^. The phasor coordinates of free NADH maps on the right side of plot with a lifetime of 0.38 ns and the protein bound form of NADH (bound with lactate dehydrogenase) maps on the left at 3.4ns (Supplementary Fig. 1b). This lifetime distribution of the free and bound forms of NADH in the phasor plot was previously described as the metabolic or M-trajectory^12,32–34^.

Next, embryos were treated with known biochemical inhibitors of oxidative phosphorylation and glycolysis^21^. Oxidative phosphorylation was inhibited at the early compaction stage with a cocktail of rotenone and antimycin A (R&A) (500nM) by inhibiting complex I and complex III of the electron transport chain. Embryos were imaged after a 4-hour culture period (Fig. 4a). The FLIM images showed increased fractional contributions of free NADH (shorter lifetimes) when compared to controls (Fig. 4a). This shift towards glycolytic metabolism is seen in a dose dependent manner (Supplementary Fig. 3), indicating that embryos cultured in R&A have decreased oxidative phosphorylation activity (Fig. 4a, b). We also cultured the early blastocyst stage embryos in 1mM 2-Deoxy-D-Glucose (2DeoxyG), an analog of glucose, to inhibit glycolysis (Fig. 4c). The glucose analog treatment shifted the phasor-FLIM distribution to longer lifetime (an increase of bound NADH) (Fig. 4c), which correlates with a decrease in glycolysis (Fig. 4c, d). These findings suggest that the source of the changes seen in the phasor coordinates throughout the pre-implantation stages in the D-trajectory is in part due to the contribution from metabolic shifts of NADH.

**Figure 4:**
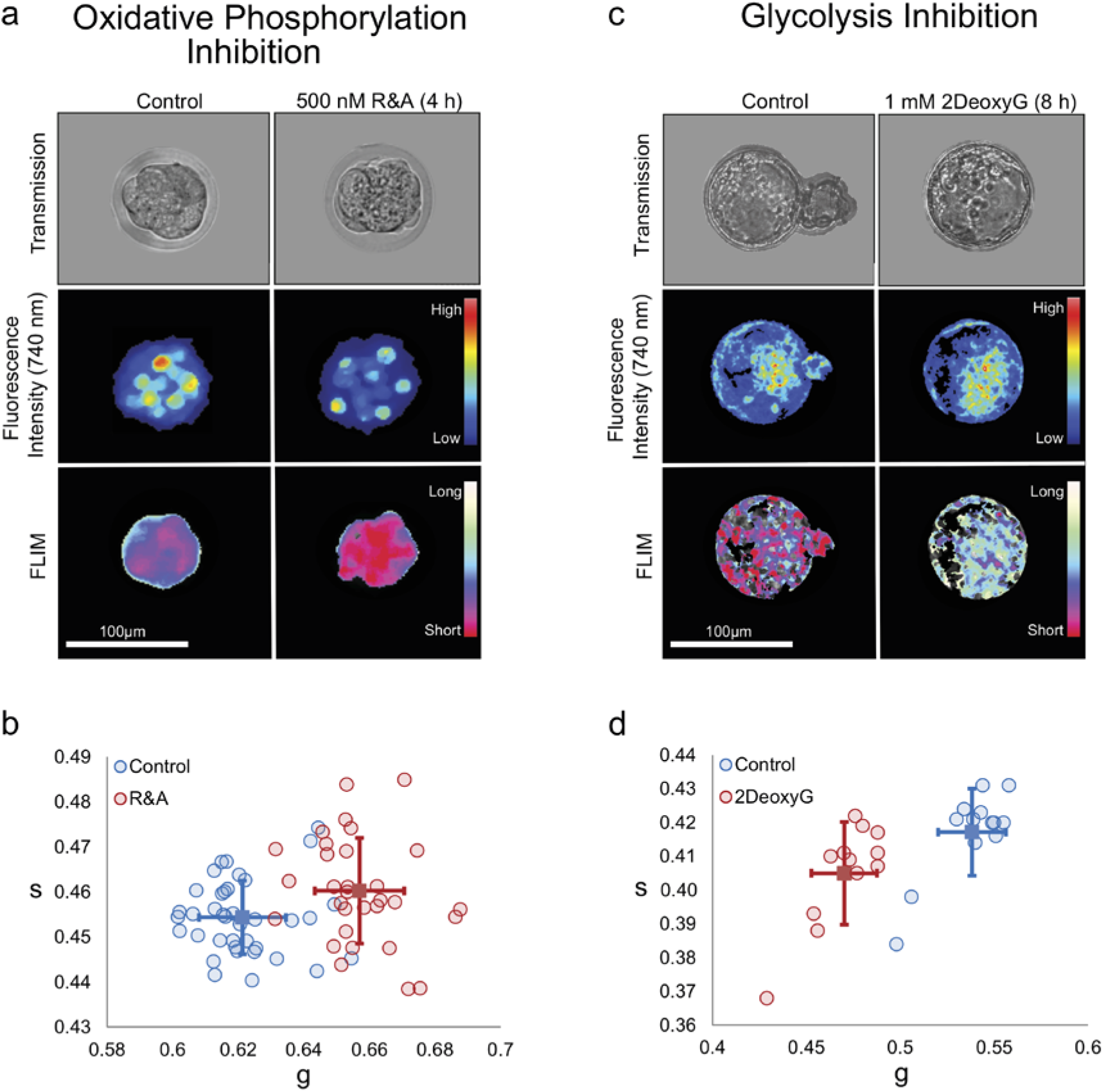
Fluorescence lifetime trajectories reveal metabolic states of pre-implantation mouse embryos. a) Transmission (top), fluorescence intensity (middle) and FLIM (bottom) images for control and 4-hour rotenone and antimycin A (R&A) treated embryos. Note a shift from long to short lifetimes (blue to red in FLIM image). b) g and s values of control and R&A-treated embryos for individual embryos. Blue circles are controls (n= 38), red circles are R&A-treated embryos (n= 31), and solid squares and the error bars in the figures means the average and variation of each group (student t-test for g value: p-value= 2.86E-16). FLIM images indicate a rightward shift from long to short lifetimes. c) Transmission (top), fluorescence intensity (middle) and FLIM (bottom) images for control and 2DeoxyG-treated embryos. Note a shift from long to short lifetimes (red to white in FLIM image). d) g and s values of control and 2DeoxyG-treated embryos. Blue squares are controls (n= 12), red circles are 2DeoxyG-treated embryos (n= 13), and the average of each group can be found in the solid colored squares (student t-test for g value: p-value= 3.88E-09). Fluorescence and FLIM images indicate a leftward shift from long to short lifetimes.

### FLIM does not disrupt embryonic development

In order to ensure the safety of the FLIM imaged embryos, we determined the optimum laser power to avoid DNA damages^35^, while allowing the rapid and robust acquisition of the FLIM signal on mouse pre-implantation embryos. We exposed 2-cell (E1.5) and morula (E2.5) stage CD1 and C57/BL6NCrl embryos to varying laser powers (1.5, 3.5, 10, and 15mW) and examined the effect on the developmental progression of embryos until the blastocyst stage^36^

(Supplementary Fig. 4a-d). In order to capture FLIM-signals of embryos taken with 1.5mW laser power, 4 times longer exposure time was required than the embryos collected at 3.5mW, 10mW, 15mW laser powers due to their low signal to noise ratio. The majority of embryos exposed to 1.5 mW and 3.5 mW laser power developed to the blastocyst and there were no significant differences between the control (non-imaged) and embryos imaged at the 2-4cell stage or morula-compaction stage, irrespective of strain differences (CD1 or C57BL/6NCrl) (Supplementary Fig. 4a-d). However, at 10mW, approximately 20% and 35% of CD1 embryos imaged at the 2-cell and compaction stages, respectively, fail to progress to the blastocysts. At 15mW, nearly 50% of CD1 and C57BL/6NCrl embryos imaged at the 2-cell stage were arrested before the compaction stage, while approximately 30% of CD1 and 12% of C57/BL6NCrl embryos imaged at the compaction stage failed to develop to blastocysts. We conclude that CD1 embryos are more sensitive to the laser damage and 3.5mW is the ideal laser power for our FLIM analysis.

Next, we examined the activation of the DNA repair pathway in the embryo by conducting immunofluorescence staining for anti-phosphorylated Histone 2AX (H2AXs139), a novel marker for DNA-double strand breaks^37,38^. Both the non-imaged and imaged embryos were indistinguishable and did not show any signs of DNA repair pathway activation at 3.5mW (Supplementary Fig. 4e). However, embryos exposed to 1.5mW laser power, which required longer laser exposure time (12 minutes, instead of ~3 minutes) showed the sign of DNA damage (Supplementary Fig. 4e). We conclude that FLIM imaging of the morula stage embryo at 3.5mW excitation is safe to use and employed for the subsequent experiments.

### FLIM distinguishes pre-implantation embryos under stress conditions

Given that early cleave stage embryos utilize aspartate, pyruvate, and lactate for energy metabolism^39^ we tested whether the unique lifetime distribution patterns of an embryo cultured under altered physiological states can be detected by the changes in spectroscopic distributions of phasor-FLIM. We cultured 2-cell and morula stage embryos in standard mouse embryo culture media (KSOMaa), flushing and holding media (FHM: DMEM-pyruvate free with HEPES), and saline solution (PBS). Brightfield images and FLIM data were collected every 4 hours over a 24-hour period (Fig. 5). At the first time-point (4 hours), the 2-cell stage embryos cultured under KSOMaa, FHM and PBS are morphologically normal (Fig. 5a, top). However, the embryos in high stress conditions (FHM and PBS) show distinct lifetime distribution patterns on the phasor-plot when compared to that of KSOMaa cultured embryos (Fig. 5b, c, Supplementary Fig 5a). Subsequently, we find that the embryos under high stress conditions fail to cleave normally and remain at the 2-cell stage, unlike KSOMaa controls (Fig. 5b, Supplementary Fig. 5a). We performed the similar analysis using morula stage embryos and found that within a few hours under high stress culture conditions, the phasor-FLIM lifetime trajectories of embryos deviate from those cultured in KSOMaa even before the embryos show any signs of abnormal morphology (Fig. 5d-f, Supplementary Fig 5b). The cell division in FHM and PBS cultured embryos also slowed down significantly (Supplementary Fig. 5b). We conclude that phasor-FLIM is a sensitive method to detect the changes in embryo metabolism upon cellular stress.

**Figure 5:**
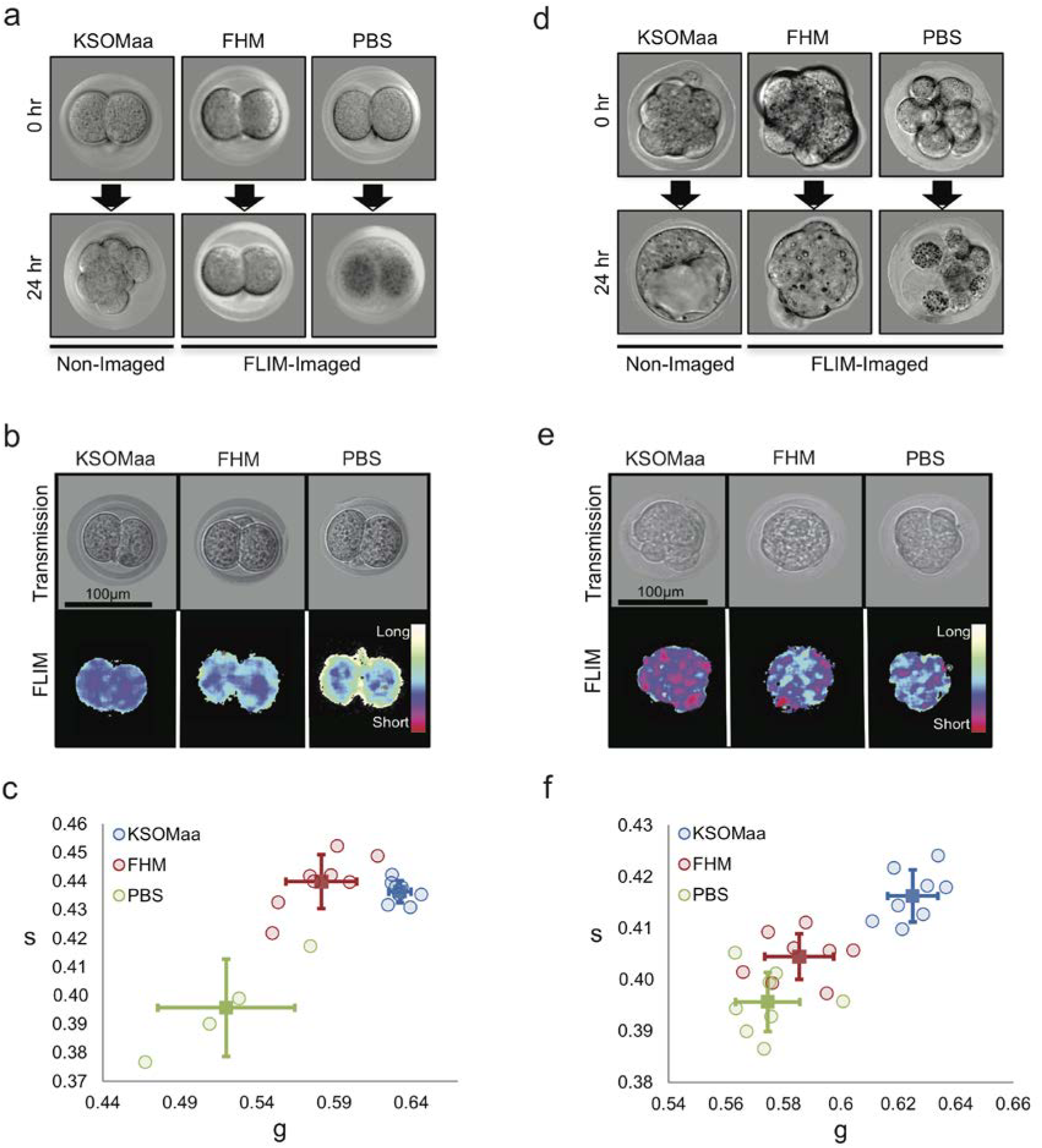
Deviation of intrinsic lifetime trajectory of embryos cultured in nutrient-depleted media. a) Transmission images of embryos collected at the 2-cell stage and cultured in KSOMaa, FHM, or PBS for 24 hours. b) Representative transmission and FLIM images of embryos in KSOMaa, FHM, or PBS for 4 hours. c) Scatter plot of g and s lifetimes collected from a group of embryos cultured in KSOMaa (n=10), FHM (n=10) and PBS (n=4) for 4 hours. p-value= 0.0002** and 0.01* (student t-test of g value) for the FHM and PBS group compare with KSOMaa group. d) Transmission images of embryos collected at the compaction stage and cultured in KSOMaa, FHM, or PBS for 24 hours. e) Representative transmission and FLIM images of embryos in KSOMaa, FHM, or PBS for 4 hours. f) Scatter plot of g and s of lifetimes collected from a group of embryos cultured in KSOMaa (n= 8), FHM (n=8), and PBS (n=8). p-value = 9.29E-06** and 3.21E-07** (student t-test of g value) for the FHM and PBS group compare with KSOMaa group.

### Derivation of the embryo viability index (EVI) for assessing the developmental potential of pre-implantation embryo

The phasor distribution analysis of pre-implantation mouse embryos allows us to distinguish between normal and highly stressed embryos (Fig. 5). Therefore, we determined whether the developmental potential of pre-implantation embryos is predictable through phasor-FLIM analysis. We first performed time-lapse phasor-FLIM imaging of embryos from the 2-cell stage for ~60 hours to identify the most desirable stage to predict the developmental potential of embryos (Fig. 6a, Supplementary Movie 2). At the end of the 60-hour culture period we classified embryos as healthy (H) if they reached the normal blastocyst stage or not healthy (NH) if embryos were arrested before reaching the blastocyst stage or displaying some abnormal morphology at the blastocyst (Fig. 6a). We then applied the distance analysis (DA) algorithm^40^ to identify key spectroscopic parameters that could differentiate healthy (H) from unhealthy (UH) embryos by machine learning. Using the DA algorithm, the 3D phasor histogram was separated into 4 sections based on the phasor coordinates (g, s) intensity, from which, 6 parameters were extracted from each section, generating a total of 24 parameters (see Methods). The healthy embryos (H group) were used as the control set and the unhealthy embryos (UH group) were used as the sample set. Each of these sets included images from multiple embryos from each stage in development. Next, we calculated the average and variance of the training set, which includes two groups (H and UH), and weighted only 20 parameters (g, s, secondary moment a, b and angle from 4 sub-layers, intensity excluded) in each set from 3D phasor plot. After optimizing the weights to maximize the difference between unhealthy and healthy group embryos, we applied these weights to index a new score called the EVI or Embryo Viability Index (Methods). This partition metric defines the degree of separation of the test embryos from the average of the training set where −1 to −10 are unhealthy embryos, and +1 to +10 are healthy embryos.

**Figure 6:**
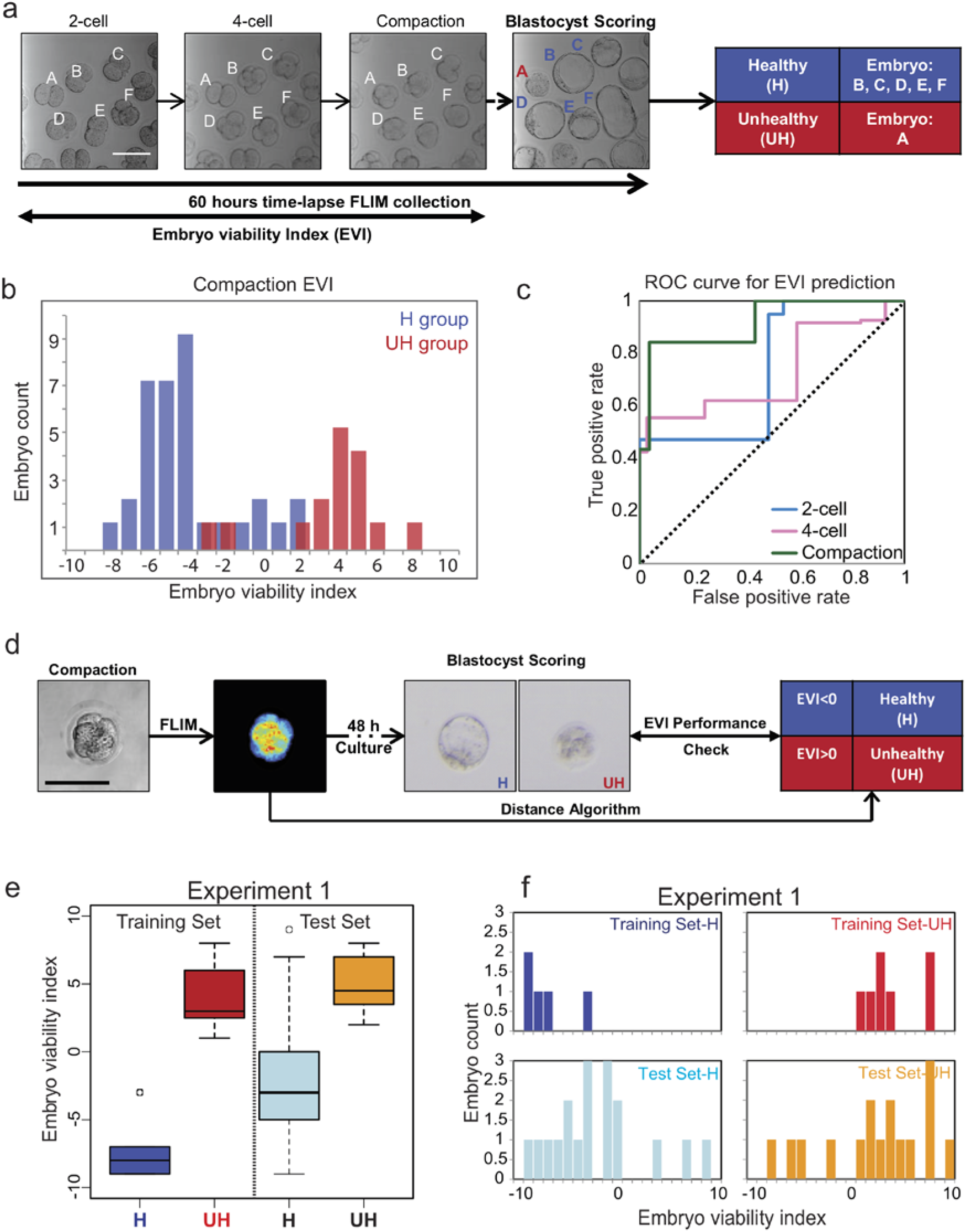
Derivation of the embryo viability index (EVI) gauging embryo quality. a) Schematic of our experimental set up. Individual embryos (A-F) were followed from the 2-cell to blastocyst stage and classified as healthy (H) and unhealthy (UH) group according to their morphology at E4.5. b) Histogram of embryo viability index (EVI) of early compaction embryos from one representative experiment (H group, n=37; UH group, n=27). The blue and red bars represent the embryo condition determined as healthy and unhealthy at ~60 hours after FLIM imaging at the pre-compaction stage. c) Receiver operating characteristic (ROC) curve shows the performance of the binary classification model developed from lifetime distribution patterns of early developmental stage embryos (2-cell, 4-cell, and early compaction stage). The area under a curve for each stage is 0.739 (2-cell), 0.728 (4-cell) and 0.916 (early compaction). The dashed line in the diagonal is presented as a random bi-classification model. d) Schematic of FLIM-Distance Analysis Pipeline. e) Box-whisker plots of experiment 1 showing training set of healthy (n=5) and unhealthy (n=7) groups and tested unknowns of healthy (n=18) and unhealthy (n=16) embryos. f) Bar graph of embryo viability index of experiment 1. Training set H is in navy, training set UH is in red. Testing set H is in light blue, and Testing set U is in orange.

Next, we examined the DA data from 2-cell, 4-cell, and the early compaction stage to determine the best binary classification model using receiver operating characteristic (ROC) curves (Fig. 6b, c, Supplementary Fig. 6a, b). We have classified the embryos predicted to be healthy in positive values (EVI<0, in blue), and embryos predicted to be unhealthy in negative values (EVI>0, in red). The plot of true positive rates against false positive rates gives an area under the ROC curve (AUC) for 2-cell, 4-cell, and the early compaction stage embryos, which were 0.739, 0.728, and 0.916, respectively. We conclude that the spectroscopic characteristics of the early compaction stage embryos (prediction accuracy with the highest AUC) possess the best parameters for separating embryos that will develop into normal blastocysts (Fig. 6b, c, Supplementary Fig. 6a, b).

An embryo viability prediction pipeline was developed based on the DA of phasor-FLIM images of the early compaction stage embryos (Fig. 6d). We have FLIM imaged embryos at the early compaction stage and all of these embryos were allowed to develop to the blastocyst equivalent stage. The resulting embryos were classified as H or UH. We then selected a small number of healthy (H) and unhealthy (UH) embryos and obtained an EVI training data set. The remaining unselected embryos were also subjected to the DA program as “unknowns” (test set) to test the predictability of EVI. In experiment 1, we followed the development of 35 morphologically healthy looking early compaction stage embryos (pooled from 4 mating pairs), until the blastocyst stage (Fig. 6e, f). Of the 34 embryos, 18 developed to normal blastocysts and thus assigned as healthy (H), and 16 embryos that failed to reach the blastocyst were assigned as unhealthy (UH). When we applied EVIs that were determined by the training set, 83.3% of healthy embryos (15 out of 18 embryos) and 75.0% of unhealthy embryos (12 out of 16 embryos) were correctly predicted by EVI (Fig. 6e, f). Subsequently, we performed another 4 biologically independent experiments using a total of 134 embryos and the results are shown in Table 1 and Supplementary Figure 6c, d. We achieved 85.9% accuracy (n=134) where a total of 88.5% healthy embryos (n=96) and 73.7% unhealthy embryos (n=38) were identified. Based on the results, we conclude that the DA program is able to predict the development potential of pre-implantation embryos at the early compaction stage.

**Table 1:** Statistical analysis of DA program as a means to predict embryo viability.

|  | False positive rate | True positive rate | Accuracy | Presicion | Sensitivity | Specificity |
| --- | --- | --- | --- | --- | --- | --- |
| *Experiment 1* | 0.250 | 0.833 | 0.794 | 0.789 | 0.833 | 0.750 |
| *Experiment 2* | 0.375 | 0.852 | 0.800 | 0.885 | 0.852 | 0.625 |
| *Experiment 3* | 0.500 | 0.920 | 0.862 | 0.920 | 0.920 | 0.500 |
| *Experiment 4* | 0.000 | 0.846 | 0.900 | 1.000 | 0.846 | 1.000 |
| *Experiment 5* | 0.333 | 1.000 | 0.938 | 0.929 | 1.000 | 0.667 |
|  | 0.292 | 0.890 | 0.859 | 0.905 | 0.890 | 0.708 |

## Discussion

Here we report that the phasor-FLIM represents a promising new approach for assessing the quality of pre-implantation mouse embryos. First, we have applied the phasor-FLIM analysis to capture developmental states during pre-implantation development. The spectroscopic trajectory, which we are calling the “D-trajectory” (D for development), is attributed to a combination of metabolic fluorescent species and production of ROS in conjunction with oxidized lipid metabolism within the embryo (Fig. 2c, d), and this trajectory correlates well with other measurements of embryonic development^41–43^. Second, the intrinsic lifetime trajectory of pre-implantation embryos cultured in nutrient-deficient media deviates from the normal lifetime distribution, indicating that the lifetime trajectory can be used to detect metabolic changes in embryos. Third, we have applied the DA program that uses spectroscopic parameters from 3D phasor histograms of embryos and shown that EVI is a non-morphological, quantitative index that can provide useful information on the quality of pre-implantation embryos.

Other spectroscopic technologies have emerged as a non-invasive means of revealing embryo viability via detection of various metabolic states of common molecules associated with embryo development. Raman, near-infrared, Nuclear Magnetic Resonance (NMR), and Fourier-transform infrared spectroscopy can also detect the metabolic states of pyruvate, lactate, glucose, and oxygen during pre-implantation mammalian development^44–46^. However, at present time these technologies suffer from a number of shortcomings. It is challenging for these approaches to analyze the data in the short time window needed for the host transfer of embryos. The data analyses are technically demanding and may not be intuitively obvious for the general clinical use. The technologies require fluid samples collected from the embryo culture media and the data are inherently noisier. Nonetheless, in the future, with improvements these spectroscopic approaches are likely to provide physicochemical parameters that will be useful in quantitating the influence of ovulation induction, oocyte retrieval, and *in vitro* culture procedures.

Development of qualitative and objective means for assessing embryo quality and viability that are safer and faster will provide significant advances in IVF and animal breeding facilities. If phasor-FLIM is to be applied for diagnostic purposes, it will be crucial to establish that the procedure does not perturb gene expression after the procedure. To date, embryos subjected to phasor-FLIM analysis appeared to be morphologically normal, and we did not detect signs of apoptosis or aberrations in nuclear morphology. However, it is possible that the phasor-FLIM procedure causes other alterations that cannot be easily detected by these morphological criteria. In the future, it will be important to perform additional molecular characterizations (i.e., DNA sequencing) to eliminate the possibility. As a means of ensuring that phasor-FLIM does not affect implantation efficacy of embryos and embryonic development, it will be also useful to determine the birth rates arising from implantation of phasor-FLIM-treated embryos and follow the developmental processes of newborns. Overall, this work has the potential to improve our understanding of energy metabolism in developing mammalian embryos and advance the ART field directly.

## Acknowledgements

We would like to thank Drs. Enrico Gratton at the Laboratory for Fluorescence Dynamics (LFD) and Katrina Waymire as well as the members of the Digman lab and the Cho lab for helpful discussion and assistance. We would also like to thank Dr. Alexander Dvornikov at the LFD for his assistance in using the DIVER microscope to measure THG. This work is supported by the NIH grants P41-RRO3155 and Hellman Fellows Fund (M.A. Digman and N. Ma) and R21HD090629 (K. Cho, M.A. Digman), grants from NSF 1562176 (K. Cho), California Institute for Regenerative Medicine 100000900 RB5-07458 (N. Ma), and the Samueli Career Development Chair (M.A, Digman), and UC President’s Dissertation Year Fellowship (N. Soledad).

## Contributions

M.A.D and S.M. initiated the phasor FLIM on embryo experiment. N.M. and S.N., M.A.D., K.C. designed the embryonic developmental tracking, metabolic pathway manipulation test, embryonic viability diagnosis test experiment. N.M. and M.A.D. designed the THG experiment. S.N., P.P. and T.Y. helped in all the mouse work and retrieving the pre-implantation embryos. N.M. did the imaging and analysis. S.N. and N.M. wrote the first draft of the manuscript, and all authors contributed to subsequent revisions.

## References

1 Baczkowski, T., Kurzawa, R. & Głabowski, W. Methods of embryo scoring in in vitro fertilization. Reprod Biol 4, 5–22 (2004).

2 De Sutter, P., Dozortsev, D., Qian, C. & Dhont, M. Oocyte morphology does not correlate with fertilization rate and embryo quality after intracytoplasmic sperm injection. Human Reproduction 11, 595–597 (1996).

3 Scott, L. A. & Smith, S. The successful use of pronuclear embryo transfers the day following oocyte retrieval. Human Reproduction 13, 1003–1013 (1998).

4 Gardner, D. K., Lane, M., Stevens, J., Schlenker, T. & Schoolcraft, W. B. Blastocyst score affects implantation and pregnancy outcome: towards a single blastocyst transfer. Fertility and sterility 73, 1155–1158 (2000).

5 Scott, R. et al. Noninvasive metabolomic profiling of human embryo culture media using Raman spectroscopy predicts embryonic reproductive potential: a prospective blinded pilot study. Fertil Steril 90, 77–83, doi:10.1016/j.fertnstert.2007.11.058 (2008).

6 O’Neill, C. & Saunders, D. Assessment of embryo quality. The Lancet 324, 1035 (1984).

7 González, R. R. et al. Leptin and leptin receptor are expressed in the human endometrium and endometrial leptin secretion is regulated by the human blastocyst 1. The Journal of Clinical Endocrinology & Metabolism 85, 4883–4888 (2000).

8 Sher, G., Keskintepe, L., Nouriani, M., Roussev, R. & Batzofin, J. Expression of sHLA-G in supernatants of individually cultured 46-h embryos: a potentially valuable indicator of ‘embryo competency’and IVF outcome. Reproductive biomedicine online 9, 74–78 (2004).

9 Botros, L., Sakkas, D. & Seli, E. Metabolomics and its application for non-invasive embryo assessment in IVF. Molecular human reproduction 14, 679–690 (2008).

10 Digman, M. A., Caiolfa, V. R., Zamai, M. & Gratton, E. The phasor approach to fluorescence lifetime imaging analysis. Biophysical journal 94, L14–L16 (2008).

11 Squirrell, J. M., Wokosin, D. L., White, J. G. & Bavister, B. D. Long-term two-photon fluorescence imaging of mammalian embryos without compromising viability. Nat Biotechnol 17, 763–767, doi:10.1038/11698 (1999).

12 Stringari, C. et al. Phasor approach to fluorescence lifetime microscopy distinguishes different metabolic states of germ cells in a live tissue. Proc Natl Acad Sci U S A 108, 13582–13587, doi:10.1073/pnas.1108161108 (2011).

13 Datta, R., Alfonso-Garcia, A., Cinco, R. & Gratton, E. Fluorescence lifetime imaging of endogenous biomarker of oxidative stress. Sci Rep 5, 9848, doi:10.1038/srep09848 (2015).

14 Watanabe, T. et al. Characterisation of the dynamic behaviour of lipid droplets in the early mouse embryo using adaptive harmonic generation microscopy. BMC Cell Biol 11, 38, doi:10.1186/1471-2121-11-38 (2010).

15 Gardner, D. K. & Wale, P. L. Analysis of metabolism to select viable human embryos for transfer. Fertil Steril 99, 1062–1072, doi:10.1016/j.fertnstert.2012.12.004 (2013).

16 Wale, P. L. & Gardner, D. K. The effects of chemical and physical factors on mammalian embryo culture and their importance for the practice of assisted human reproduction. Hum Reprod Update 22, 2–22, doi:10.1093/humupd/dmv034 (2016).

17 Leese, H. J. & Barton, A. M. Pyruvate and glucose uptake by mouse ova and preimplantation embryos. J Reprod Fertil 72, 9–13 (1984).

18 Dumollard, R., Carroll, J., Duchen, M., Campbell, K. & Swann, K. in Seminars in cell & developmental biology. 346–353 (Elsevier).

19 Wright, B. K. et al. Phasor-FLIM analysis of NADH distribution and localization in the nucleus of live progenitor myoblast cells. Microsc Res Tech 75, 1717–1722, doi:10.1002/jemt.22121 (2012).

20 Stringari, C., Nourse, J. L., Flanagan, L. A. & Gratton, E. Phasor fluorescence lifetime microscopy of free and protein-bound NADH reveals neural stem cell differentiation potential. PLoS One 7, e48014, doi:10.1371/journal.pone.0048014 (2012).

21 Stringari, C. et al. Metabolic trajectory of cellular differentiation in small intestine by Phasor Fluorescence Lifetime Microscopy of NADH. Sci Rep 2, 568, doi:10.1038/srep00568 (2012).

22 Stringari, C., Sierra, R., Donovan, P. J. & Gratton, E. Label-free separation of human embryonic stem cells and their differentiating progenies by phasor fluorescence lifetime microscopy. J Biomed Opt 17, 046012, doi:10.1117/1.JBO.17.4.046012 (2012).

23 Shyh-Chang, N., Daley, G. Q. & Cantley, L. C. Stem cell metabolism in tissue development and aging. Development 140, 2535–2547, doi:10.1242/dev.091777 (2013).

24 Covarrubias, L., Hernández-García, D., Schnabel, D., Salas-Vidal, E. & Castro-Obregón, S. Function of reactive oxygen species during animal development: passive or active? Developmental biology 320, 1–11 (2008).

25 Ray, P. D., Huang, B.-W. & Tsuji, Y. Reactive oxygen species (ROS) homeostasis and redox regulation in cellular signaling. Cellular signalling 24, 981–990 (2012).

26 Esterbauer, H. Cytotoxicity and genotoxicity of lipid-oxidation products. Am J Clin Nutr 57, 779S–785S; discussion 785S-786S (1993).

27 Porter, N. A., Wolf, R. A. & Weenen, H. The free radical oxidation of polyunsaturated lecithins. Lipids 15, 163–167, doi:10.1007/bf02540963 (1980).

28 Gardner, D. K., Lane, M., Calderon, I. & Leeton, J. Environment of the preimplantation human embryo in vivo: metabolite analysis of oviduct and uterine fluids and metabolism of cumulus cells. Fertil Steril 65, 349–353 (1996).

29 Lane, M. & Gardner, D. K. Lactate Regulates Pyruvate Uptake and Metabolism in the PreimplantationMouse Embryo. Biology of reproduction 62, 16–22 (2000).

30 Chance, B., Cohen, P., Jobsis, F. & Schoener, B. Intracellular oxidation-reduction states in vivo. Science 137, 499–508 (1962).

31 Hamaoka, T., McCully, K. K., Quaresima, V., Yamamoto, K. & Chance, B. Near-infrared spectroscopy/imaging for monitoring muscle oxygenation and oxidative metabolism in healthy and diseased humans. J Biomed Opt 12, 062105, doi:10.1117/1.2805437 (2007).

32 Skala, M. C. et al. In vivo multiphoton microscopy of NADH and FAD redox states, fluorescence lifetimes, and cellular morphology in precancerous epithelia. Proc Natl Acad Sci U S A 104, 19494–19499, doi:10.1073/pnas.0708425104 (2007).

33 Bird, D. K. et al. Metabolic mapping of MCF10A human breast cells via multiphoton fluorescence lifetime imaging of the coenzyme NADH. Cancer Res 65, 8766–8773, doi:10.1158/0008-5472.CAN-04-3922 (2005).

34 Wu, D. et al. Metabolic complementarity and genomics of the dual bacterial symbiosis of sharpshooters. PLoS Biol 4, e188, doi:10.1371/journal.pbio.0040188 (2006).

35 Wang, Z. W. et al. Laser microbeam-induced DNA damage inhibits cell division in fertilized eggs and early embryos. Cell Cycle 12, 3336–3344, doi:10.4161/cc.26327 (2013).

36 Cross, J. C., Werb, Z. & Fisher, S. J. Implantation and the placenta: key pieces of the development puzzle. Science 266, 1508–1518 (1994).

37 Revet, I. et al. Functional relevance of the histone gammaH2Ax in the response to DNA damaging agents. Proc Natl Acad Sci U S A 108, 8663–8667, doi:10.1073/pnas.1105866108 (2011).

38 Sonoda, E. et al. Collaborative roles of gammaH2AX and the Rad51 paralog Xrcc3 in homologous recombinational repair. DNA Repair (Amst) 6, 280–292, doi:10.1016/j.dnarep.2006.10.025 (2007).

39 Lane, M. & Gardner, D. K. Mitochondrial malate-aspartate shuttle regulates mouse embryo nutrient consumption. Journal of Biological Chemistry 280, 18361–18367 (2005).

40 Ranjit, S., Dvornikov, A., Levi, M., Furgeson, S. & Gratton, E. Characterizing fibrosis in UUO mice model using multiparametric analysis of phasor distribution from FLIM images. Biomedical Optics Express 7, 3519–3530 (2016).

41 Biggers, J., Whittingham, D. & Donahue, R. The pattern of energy metabolism in the mouse oöcyte and zygote. Proceedings of the National Academy of Sciences 58, 560–567 (1967).

42 Brinster, R. L. Studies on the development of mouse embryos in vitro. Journal of reproduction and fertility 10, 227–240 (1965).

43 Whitten, W. K. Culture of tubal ova. Nature 179, 1081–1082 (1957).

44 Seli, E. et al. Noninvasive metabolomic profiling of embryo culture media using Raman and near-infrared spectroscopy correlates with reproductive potential of embryos in women undergoing in vitro fertilization. Fertil Steril 88, 1350–1357, doi:10.1016/j.fertnstert.2007.07.1390 (2007).

45 Vergouw, C. G. et al. Metabolomic profiling by near-infrared spectroscopy as a tool to assess embryo viability: a novel, non-invasive method for embryo selection. Hum Reprod 23, 1499–1504, doi:10.1093/humrep/den111 (2008).

46 Seli, E. et al. Noninvasive metabolomic profiling as an adjunct to morphology for noninvasive embryo assessment in women undergoing single embryo transfer. Fertil Steril 94, 535–542, doi:10.1016/j.fertnstert.2009.03.078 (2010).

47 König, K., So, P. C., Mantulin, W., Tromberg, B. & Gratton, E. Two - photon excited lifetime imaging of autofluorescence in cells during UV A and NIR photostress. Journal of microscopy 183, 197–204 (1996).

48 *ISS | Technical Notes | Fluorescence Lifetime*, <http://www.iss.com/resources/research/technical_notes/K2CH_FLT.html> (2016).

49 Chiang, M. et al. Analysis of in vivo single cell behavior by high throughput, human-in-the-loop segmentation of three-dimensional images. BMC Bioinformatics 16, 397, doi:10.1186/s12859-015-0814-7 (2015).

50 Kukreti, S. et al. Characterization of metabolic differences between benign and malignant tumors: high-spectral-resolution diffuse optical spectroscopy. Radiology 254, 277–284 (2009).

