## Supplementary Materials for "Label-free assessment of pre-implantation embryo quality by the Fluorescence Lifetime Imaging Microscopy (FLIM)-phasor approach" (from another version of this preprint: version 1, 2018-03-22)

**Supplementary Figure Legends:**

*Supplementary Figure 1: Phasor FLIM analysis.* a) Phasor FLIM analysis. During FLIM collection, a pulsed 2-photon laser is used to measure the intensity at short time windows (time arrival of the photons) as a function of time. Instead of fitting the decay curve into an exponential equation (black line), the raw data (intensity at each pixel) is transformed into polar coordinates by plotting the sine (red line) and cosine (blue line) using Fourier transformation, for every pixel in the object, the fluorescence lifetime can be obtained as “phasor lifetime” b) Phasor fingerprint of pure intrinsic biomarkers of free NADH in solution, bound NADH in the presence of lactate dehydrogenase, and a long lifetime species derived from lipid droplets. Given that the free form of NADH exhibits a compact structure with a low fluorescence quantum yield (ϕ=0.019) and a short lifetime of 0.4ns and the extended form of NADH bound to lactate dehydrogenase with a much higher quantum yield (ϕ= 0.099) with a longer fluorescence lifetime up to ~3.4ns, the lifetimes of these two states can be easily distinguished^7,8^. Based on the law of phasor addition, any sample contains the combination signature of these three species will fall within the triangle joining the three phasors

*Supplementary Figure 2:* *D-trajectory is composed of two distinct trajectories that correlate with embryo stages.* a) Representative phasor plots for the pre-implantation mouse embryo from early cleavage stage to blastocyst stage. b) *Detection of reactive oxygen species in pre-implantation mouse embryos.* 2', 7’-dichlorodihydrofluorescein diacetate (DCF-DA) stain across pre-implantation mouse embryos. c) Four examples the D-Trajectory observed throughout pre-implantation stages (E1.5-E4.0).

*Supplementary Figure 3: Fluorescence lifetime trajectories reveal metabolic states changes of pre-implantation mouse embryos and shows dose dependent manner.* a) Transmission (top), fluorescence (middle) and FLIM (bottom) images for control and 4-hour 100nM and 500nM rotenone and antimycin A (R & A) treated embryos, indicating a shift from long to short lifetimes. b) g and s values of control and 4-hour 100nM and 500nM R&A-treated embryos for individual embryos. Blue circles are controls (n= 38), red circles are 4-hour 100nM R&A-treated embryos (n= 21), and green circles are 4-hour 500nM R&A-treated embryos (n=31). And the average of each group can be found in the solid blue and red circles (for g value of 100nM treatment group and 500nM group compare with control group, p-value=1.76E-5, and 2.86E-16, respectively). FLIM images indicate a rightward shift from long to short lifetimes. Student T-Test, two tailed tests.

*Supplementary Figure 4a, b: Validating the safety of FLIM-imaging of E1.5 pre-implantation mouse embryos.* a) Transmission images of embryos before and after imaging and culturing for 72 hours till E4.5. b) Assessment of embryonic development after imaging and 72-hour *in vitro* culture reported as percent development. For CD1: E1.5 non-image control (n=72); 1.5mW(n=42); 3.5mW(n=42), 10mW(n=31), 15mW(n=45). For C57BL/6NCrl: E1.5 non-image control (n=37); 1.5mW(n=37); 3.5mW(n=36), 10mW(n=38), 15mW(n=37). Image size is 708.49*708.49 µm.

*Supplementary Figure 4c, d: Validating the safety of FLIM-imaging of E2.5 pre-implantation mouse embryos.* c) Transmission images of embryos before and after imaging and culturing for 48 hours till E4.5. d) Assessment of embryonic development after imaging and 48-hour *in vitro* culture reported as percent development. For CD1: E2.5 non-image control (n=53); 1.5mW(n=43); 3.5mW(n=43), 10mW(n=28), 15mW (n=28, p<0.05 *). For C57/BL6NCrl: E2.5 non-image control (n=75); 1.5mW(n=42); 3.5mW(n=47), 10mW(n=44), 15mW (n=41 p<0.001 ***). Student T-Test, one tailed test. Everything is nonsignificant except the ones annotated. Image size is 708.49*708.49 µm.

*Supplementary Figure 4e: Validating the safety of FLIM-imaging of pre-implantation mouse embryos by H2AX staining for Assessment of DNA Damage response initiation of DNA repair pathways.* Hoechst (blue) and H2AX(red) stain of E2.5 non-imaged and imaged embryos. For CD1: E2.5 control (n=3); 1.5mW(n=3); 3.5mW(n=3), 10mW(n=3), 15mW(n=3)

For C57/BL6NCrl: E2.5 control (n=9); 1.5mW(n=9); 3.5mW(n=8), 10mW(n=6), 15mW(n=8). Scale bar set to 20µm.

*Supplementary Figure 5: Average number of cells per embryos cultured under high stress conditions.* a) Bar graph with the average number of cells per embryo after the culturing period for embryos starting at the 2-cell stage (KSOMaa non-image control (n=8), KSOMaa (n=10), FHM (n=11), PBS (n=11), p-value = 0.004** and 0.001**). b) Bar graph with the average number of cells per embryo after the culturing period for embryos starting at the morula stage (KSOMaa non-image control (n=10), KSOMaa (n=18), FHM (n=11), PBS (n=11), p-value = 0.002** and 0.02*).

*Supplementary Figure 6 Part 1: Embryo viability index of morula shows the potential to distinguish healthy and unhealthy pre-cleavage stage embryos.* a) Histogram of embryo viability index of 2-cell and morula embryos from one representative experiment (2-cell, H group, (n=18), UH group, (n=17). 4-cell, H group, (n=25), UH group, (n=9)). The blue and red bars represent the morula stage FLIM-fingerprints of healthy (H) and unhealthy (UH) embryos after image ~60 hours, respectively. b) Receiver operating characteristic (ROC) curve shows the performance of the binary classification model developed from lifetime distribution patterns of pre-compaction stage embryos (2-, 4-, and early compaction) of two time-lapse FLIM tracking experiment. The area under curve for each stage is 0.777 (2-cell, H, n=37, UH, n=8), 0.823 (4-cell, H, n=45, UH, n=8) and 1.000 (early compaction, H, n=30, UH, n=2) for experiment 2 and 0.777 (2-cell, H, n=38, UH, n=10), 0.813 (4-cell, H, n=39, UH, n=7) and 0.945 (early compaction, H, n=39, UH, n=6) for experiment 3.

*Supplementary Figure 6 Part 2: Embryo viability index predicts healthy and unhealthy embryos from 4 experiments.* a) Box-whisker plot showing training set of healthy (H) and unhealthy (UH) groups and tested unknowns of healthy (H) and unhealthy (UH) embryos for experiment 2, 3, 4, and 5. Training set H are in navy, Training set UH is in red, Healthy in light blue, and Unhealthy in orange.

The n number for the training set healthy group was n= 5, 11, 2, 3. The n number for the training set unhealthy group was n= 4, 3, 2, 2. The n number for the test unknown healthy group was n= 27, 25, 13, 13. The n number for the test unknown unhealthy group was n= 8, 4, 7, 3. b) Bar graph of embryo viability index of experiment 2, 3, 4 and 5. Training set H are in navy, Training set UH is in red, Healthy in light blue, and Unhealthy in orange.

**Supplementary Movie 1: 3D THG images of the representative embryos from different pre-implantation embryos. a) 2-cell, b) 8-cell, c) Compaction, d) Early Blastocyst, e) Blastocyst**

**Supplementary Movie 2: 60-hour time-lapse imaging of pre-implantation embryo development from E1.5 to E4.0.** Embryos from H group (blue) are the representative healthy embryos that developed to the blastocyst stage after 60 hours of time-lapse imaging. Embryos from UH group (red) is an embryo arrested at late compaction stage after 60 hours.
